# Standardisation Challenges in ELISA-Based Quantification of Equine Chorionic Gonadotropin (eCG/PMSG)

**DOI:** 10.64898/2026.10.29.762762

**Authors:** Polina Mishchenko, Ricardo Matiz, Thorsten Buch, Martina Lösle

## Abstract

Equine chorionic gonadotropin (eCG/PMSG) is widely used for female-cycle synchronization and estrus induction in livestock and in biomedical research, where reliable quantitative assessment is increasingly important. However, its structural and glycosylation heterogeneity complicates antibody-based measurement and assay standardization. This Technical Note evaluated six commercially available quantitative PMSG ELISA kits representing three distinct analytical profile groups. Kit-provided calibration standards were characterized by Western blotting, Coomassie staining, and LC–MS/MS, while quantitative performance was assessed using serial dilutions of two commercial PMSG preparations. DRG, Creative Diagnostics, and Demeditec showed similar calibration-standard profiles and concentration-dependent responses, although measured concentrations were consistently below nominal values. Biomatik and MyBiosource displayed closely related analytical profiles but little meaningful concentration-dependent response, whereas Krishgen measurements remained within a narrow range despite changes in nominal PMSG input. The calibration standards differed substantially in protein composition, and no eCG-derived peptides were identified by LC–MS/MS under the conditions used. These findings demonstrate that quantitative results obtained with commercially available PMSG ELISAs are not necessarily comparable and may be strongly influenced by calibration-material composition, assay format, and matrix effects. The findings highlight the limitations of current quantitative PMSG assays and the need for a well-characterized, commutable reference material, together with assay-specific validation of dilution parallelism and analyte recovery, to improve the reliability and comparability of eCG/PMSG measurements.

## Introduction

Pregnant Mare Serum Gonadotropin (PMSG), also known as Equine Chorionic Gonadotropin (eCG), is a glycoprotein hormone secreted by pregnant mares and released into the maternal blood-stream during early to mid-pregnancy. Owing to its dual follicle-stimulating hormone (FSH) and luteinizing hormone (LH) activities, PMSG is widely used in reproductive management, particularly in livestock breeding [1]. It is commonly applied to induce estrus and ovulation, synchronize reproductive cycles, and promote superovulation in assisted reproduction programs. Its prolonged biological half-life, largely attributed to extensive glycosylation, further enhances its effectiveness by enabling sustained ovarian stimulation [1]. From an analytical perspective, PMSG represents a particularly challenging target. The hormone is a heterodimeric glycoprotein composed of a common to glycoprotein hormones family α-subunit and a hormone-specific β-subunit [2, 3] that are non-covalently associated and stabilized by the conserved disulfide-rich cystine-knot architecture [3]. PMSG is extensively glycosylated [4, 5], including N-linked glycans on both subunits and O-linked glycans in the extended C-terminal region of the β-subunit, with additional heterogeneity arising from differences in branching, sialylation, core fucosylation, and sialic-acid modifications [6]. This structural heterogeneity may influence both biological activity [7] and molecular recognition, resulting in differences in receptor interaction as well as antibody binding. The biological activity of PMSG has historically been assessed using *in vivo* bioassays, most notably the Cole and Erway ovarian weight gain assay [8] or Combarnous et al. modification of this assay [9], which can provide an integrated readout of receptor-mediated signaling and physiological response. However, these assays are designed to evaluate bioactivity rather than hormone amounts and are not suitable for determining the concentration of PMSG in complex samples. In contrast, enzyme-linked immunosorbent assays (ELISAs) are widely used for the detection and quantification of PMSG based on antibody recognition of specific epitopes. ELISAs enable sensitive and operationally convenient assessment of hormone presence and are commonly applied in both veterinary diagnostics and research laboratories. Importantly, ELISA-based measurements are independent of receptor binding and biological activity and are therefore intended to report immunoreactive PMSG rather than functional potency [10]. For PMSG, reliable interpretation of ELISA results is complicated by the hormone’s intrinsic structural heterogeneity [2]. Variations in glycosylation and molecular composition affect epitope accessibility and antibody affinity, thereby influencing assay sensitivity, linearity, and comparability across kits and laboratories [11]. Quantitative PMSG measurements are used to compare commercial and experimental preparations, yet the comparability of concentrations obtained with different ELISA kits has not been established. The historical WHO International Standard for equine serum gonadotrophin (NIBSC code 62/001 [12]) was established for bioassay calibration. However, stocks of reference material are no longer commercially available, and the European Pharmacopoeia Commission has suspended the monograph on equine serum gonadotrophin for veterinary use as of Ph. Eur. Supplement 10.8. In the absence of widely accepted standardized reference materials, the analytical performance of commercially available PMSG ELISA kits cannot be assumed a priori and must be evaluated in the context of both assay design and target protein properties. ELISA results may be interpreted qualitatively, semi-quantitatively, or quantitatively, depending on the assay design and calibration [10]. Qualitative assays indicate whether the target is present, while semi-quantitative assays compare relative signal between samples. Quantitative assays estimate analyte concentration from a calibration curve generated using standards with assigned values. However, the presence of a standard curve does not by itself ensure reliable quantification; the response must remain proportional across the assay range, and the calibrator must behave comparably to the tested samples. For quantitative protein analysis, ELISA is commonly implemented in direct, indirect, competitive, and sandwich formats. For PMSG/eCG, competitive and especially sandwich assays are most relevant because they are better suited to heterogeneous analytes and low-abundance targets. Most commercial PMSG/eCG kits use sandwich ELISA, in which two antibodies recognize distinct epitopes, often with secondary-antibody amplification to increase sensitivity. However, due to PMSG heterogeneity, epitope accessibility and antibody affinity can vary, which may compromise linearity, proportionality to nominal concentration, and inter-kit comparability [11]. For our experimental work, commercially available PMSG preparations are routinely used as positive controls, as they are supplied at defined nominal concentrations and are commonly assumed to provide a reliable reference. In this context, we primarily employed two widely used commercial preparations, Folligon® PMSG (MSD Animal Health; 1000 IU) and Oppa (Aviva Systems Biology; 1000 IU).

Variability in biological activity between different PMSG products is well recognized and can arise from batch-to-batch differences in production [13], such variability may also substantially affect antibody-based detection by ELISA. ELISA kits typically include calibration standards with defined PMSG concentrations to enable quantitative normalization. However, during routine measurements, ELISA-based quantification of commercial preparation Folligon showed PMSG concentrations that deviated markedly from the nominal values provided by the manufacturer.

Although the underlying cause could not be determined at this stage, this observation suggested limitations in the quantitative interpretability of PMSG ELISA measurements. In this technical note, we systematically evaluate commercially available PMSG ELISA kits with respect to their analytical consistency, quantitative performance, and practical scope. By comparing assay outputs and underlying technological approaches, we aim to identify which assay designs are appropriate for qualitative detection and which enable more robust quantitative assessment of immunoreactive PMSG. Several companies offer ELISA kits designed for the quantitative detection of PMSG in biological samples. These kits claim to provide high specificity and accuracy in detecting PMSG levels, using monoclonal antibodies targeting a unique antigenic site on the PMSG molecule (shown in Table 1)

**Table 1:** Comparison of commercially available PMSG ELISA kits.

| PMSG ELISA kit Comparison |  |  |  |  |  |  |
| --- | --- | --- | --- | --- | --- | --- |
|  | **PMSG ELISA**<br>**DRG**<br>**instruments**<br>**Germany** | **PMSG ELISA**<br>**CD Creative**<br>**Diagnostics** | **PMSG ELISA**<br>**Demeditec**<br>**Diagnostics**<br>**Germany** | **Pregnant Mare**<br>**Serum**<br>**Gonadotropin**<br>**ELISA kit My**<br>**Biosource** | **Pregnant Mare**<br>**Serum**<br>**Gonadotropin**<br>**ELISA kit**<br>**Biomatik** | **GenLISA**<br>**PMSG ELISA**<br>**Krishgen**<br>**Biosystems** |
| Quantitative ELISA | Yes | Yes | Yes | Yes | Yes | Yes |
| Type of ELISA | Sandwich | Sandwich | Sandwich | Competitive | Competitive | Sandwich |
| Antibody Specificity | 1.Monoclonal AB against PMSG epitope 2.Anti PMSG-AB-HRP | 1.Monoclonal AB against PMSG epitope 2.Anti PMSG-AB-HRP | 1.Monoclonal AB against PMSG epitope 2.Anti PMSG-AB-HRP | 1.Goat anti-rabbit AB 2.Anti-PMSG-AB-HRP | 1.Goat anti-rabbit AB 2. Anti-PMSG-AB-HRP | 1.Monoclonal anti-PMSG 2.Monoclonal Biotin-labeled anti-PMSG-AB 3.Streptavidin -HRP |
| Cross-reactions with other molecules | LH human 0.00%, FSH human 9.06%, Prolactin 0.00%, β-hCG 0.06%, hCG human 0.00% | LH human 0.00%, FSH human 9.06%, Prolactin 0.00%, β-hCG 0.06%, hCG human 0.00% | LH human 0.00%, FSH human 9.06%, Prolactin 0.00%, β-hCG 0.06%, hCG human 0.00% | No known cross-reaction factors except chemical lysis buffers | No known cross-reaction factors except chemical lysis buffers | Sodium azide |
| Assay range [mIU/mL] | 2 - 800 | 2 - 800 | 2 - 800 | 5 - 2000 | 5 - 2000 | 100 - 1600 |
| Limits and sensitivity [mIU/mL] | Limit of Blank: 0.89<br>Limit of Detection: 2.02<br>Limit of Quantification: 5.92 | Limit of Blank: 0.89<br>Limit of Detection: 2.02<br>Limit of Quantification: 5.92 | Limit of Blank: 0.89<br>Limit of Detection: 2.02<br>Limit of Quantification: 5.92 | Lower Limit of Detection: 6 | Lower Limit of Detection: 6 | Lower Limit of Quantification: 98.5 |
| Precision (CV) | Intra: 4 - 6%<br>Inter: 6.4 -8.8%<br>Inter lot: 3-10.5% | Intra: 4 - 6%<br>Inter: 6.4 -8.8%<br>Inter lot: 3-10.5% | Intra: 4 - 6%<br>Inter: 6.4 -8.8%<br>Inter lot: 3-10.5% | Intra: < 15%<br>Inter: < 15% | Intra: < 15%<br>Inter: < 15% | Intra: < 15%<br>Inter: < 18% |
| Sample type | Not specified, serum samples should be diluted ( 1:10 or higher) to avoid cross reaction with FSH | Not specified, serum samples should be diluted ( 1:10 or higher) to avoid cross reaction with FSH | Not specified, serum samples should be diluted ( 1:10 or higher) to avoid cross reaction with FSH | Plasma | Plasma | Serum, plasma, cell culture supernatant and other biological samples. Validated for citrated/EDTA plasma only. |
| Sample volume | 50 μL | 50 μL | 50 μL | 50 μL | 50 μL | 40 μL |
| Number of steps in protocol | 12 | 12 | 12 | 9 | 9 | 12 |
| Incubation times | 60 min + 60 min + 30 min = 2.5h | 60 min + 60 min + 30 min = 2.5h | 60 min + 60 min + 30 min = 2.5h | 60 min + 15 min = 75 mins | 60 min + 15 min = 75 mins | 60 min + 10 min = 70 min |
| Wash requirements | Manual washing or plate washer; washing procedure influences sensitivity and precision of the assay | Manual washing or plate washer; washing procedure influences sensitivity and precision of the assay | Manual washing or plate washer; washing procedure influences sensitivity and precision of the assay | Manual wash or plate washer for higher precision: 1x TBS | Manual wash or plate washer for higher precision: 1x TBS | Manual with wash buffer or plate washer |

| Continuation of Table 1 |  |  |  |  |  |  |
| --- | --- | --- | --- | --- | --- | --- |
|  | **PMSG ELISA**<br>**DRG**<br>**instruments**<br>**Germany** | **PMSG ELISA**<br>**CD Creative**<br>**Diagnostics** | **PMSG ELISA**<br>**Demeditec**<br>**Diagnostics**<br>**Germany** | **Pregnant Mare**<br>**Serum**<br>**Gonadotropin**<br>**ELISA kit My**<br>**Biosource** | **Pregnant Mare**<br>**Serum**<br>**Gonadotropin**<br>**ELISA kit**<br>**Biomatik** | **GenLISA**<br>**PMSG ELISA**<br>**Krishgen**<br>**Biosystems** |
| Calculations | four-parameter<br>Logistics is the<br>preferred method.<br>Linear standard<br>curve also<br>possible | four-parameter<br>Logistics is the<br>preferred method.<br>Linear standard<br>curve also<br>possible | four-parameter<br>Logistics is the<br>preferred method.<br>Linear standard<br>curve also<br>possible | “Curve Expert”<br>for standard.<br>Calculates the<br>standard curve<br>via 4 parameter<br>logistic curve fit.<br>Linear standard<br>curve also<br>possible. | “Curve Expert”<br>for standard.<br>Calculates the<br>standard curve<br>via 4 parameter<br>logistic curve fit.<br>Linear standard<br>curve also<br>possible. | Use the standard<br>calibration curve<br>method. For<br>automated<br>analysis, use<br>software with a<br>cubic spline or<br>4-PL curve fit. |

## Materials and Methods

### Materials

#### PMSG

Pregnant mare serum gonadotropin (PMSG) was obtained as Folligon® Lot: A239A02 (MSD Animal Health, Boxmeer, The Netherlands) and PMSG Protein (OPPA01037) Lot 917PGSMP07 (Aviva Systems Biology, San Diego, USA), both supplied as 1000 IU per vial. The preparation was each reconstituted according to the manufacturer’s instructions to a final concentration of 1000 IU/mL.

#### Antibody anti-PMSG

Polyclonal anti-PMSG antibody from OriGene, AP02036SU-N Lot # AK-303E110872Z-I64

### Methods

#### ELISA

Protein samples were analysed for PMSG concentrations using the respective ELISA according to the manufacturer’s instructions. Briefly, 50 µL of standards, controls, and diluted samples were added to microtiter wells pre-coated with a monoclonal antibody specific to PMSG. After a 60-minute incubation at room temperature, wells were washed three times with wash buffer. Subsequently, 100 µL of enzyme conjugate was added to each well, followed by another 60-minute incubation at room temperature. Following three additional washes, 100 µL of substrate solution was added, and the plate was incubated in the dark for 30 minutes at room temperature. The reaction was terminated by adding 50 µL of stop solution, and absorbance was measured at 450 nm using a microplate reader THE SPARK® from Tecan. PMSG concentrations were determined by constructing a standard curve from the provided calibrators. This information applies to most of the kits tested and provides a general overview of the procedure. A detailed overview and direct comparison of the individual protocols are provided in Supplementary Table 1: Overview of used ELISA kits and protocol comparison.

**Table 2:** PMSG ELISA kits order information.

| PMSG ELISA kit information |  |  |  |  |  |  |
| --- | --- | --- | --- | --- | --- | --- |
|  | PMSG ELISA<br>DRG in-<br>struments<br>Germany | PMSG ELISA<br>CD Creative<br>Diagnostics | PMSG ELISA<br>Demeditec Di-<br>agnostics Ger-<br>many | Pregnant<br>Mare Serum<br>Gonadotropin<br>ELISA kit My<br>Biosource | Pregnant<br>Mare Serum<br>Gonadotropin<br>ELISA kit<br>Biomatik | GenLISA<br>PMSG ELISA<br>Krishgen<br>Biosystems |
| Lot # | 17K095-2 | 17K104 | 17K055 | C1465081439 | C1478041497 | EPMSG0425 |
| Ref. / Cat.# | EIA-1298 | DEIA2235 | DE1298 | MBS701410 | EKC38379 | KLV0061 |

#### Standard curve calculation

For standard-curve fitting and concentration calculations, a 4PL method was used. For the method description and Python code, see the link below.

https://github.com/prostopolinushka/4PL-ELISA-analysis.git

#### Statistics analysis

For the comparison of nominal and ELISA-measured PMSG concentration technical replicates were first averaged within each independent ELISA experiment. For each ELISA kit, PMSG preparation, and nominal concentration, results are presented as the mean ± standard deviation (SD) across independent experimental runs. Mean measured concentrations were analysed by ordinary least-squares linear regression against the corresponding nominal concentrations. Regression slope, intercept, and coefficient of determination (*R*^2^) were calculated. The identity line, defined by a slope of 1 and an intercept of 0, was used as the theoretical reference for perfect agreement between nominal and measured concentrations. Deviation from nominal concentrations was additionally expressed as the absolute relative deviation and averaged across the tested concentration range. Because the evaluated ELISA kits differed in assay format and calibration characteristics, no formal hypothesis testing or pairwise statistical comparison between manufacturers was performed. Comparisons between assays are therefore descriptive.

### Western Blot

#### Sample preparation

For comparative analysis, the highest standard concentration from each ELISA kit was serially diluted to a final concentration of 4 mIU/mL prior to electrophoresis (Figure 1). A 0.9% NaCl solution was used as the dilution solvent. All samples were prepared under non-reducing conditions. For Figure 2, the kit standard samples from DRG, Creative Diagnostics, and Demeditec were diluted 1:50 to achieve optimal visualization. This dilution was not necessary for the Biomatik, MyBiosource, and Krishgen samples because of their lower overall protein content. A 0.9% NaCl solution was used as the solvent. For comparison, a commercial protein (Folligon) was used as a positive control at 1000 IU/mL.

**Figure 1:**
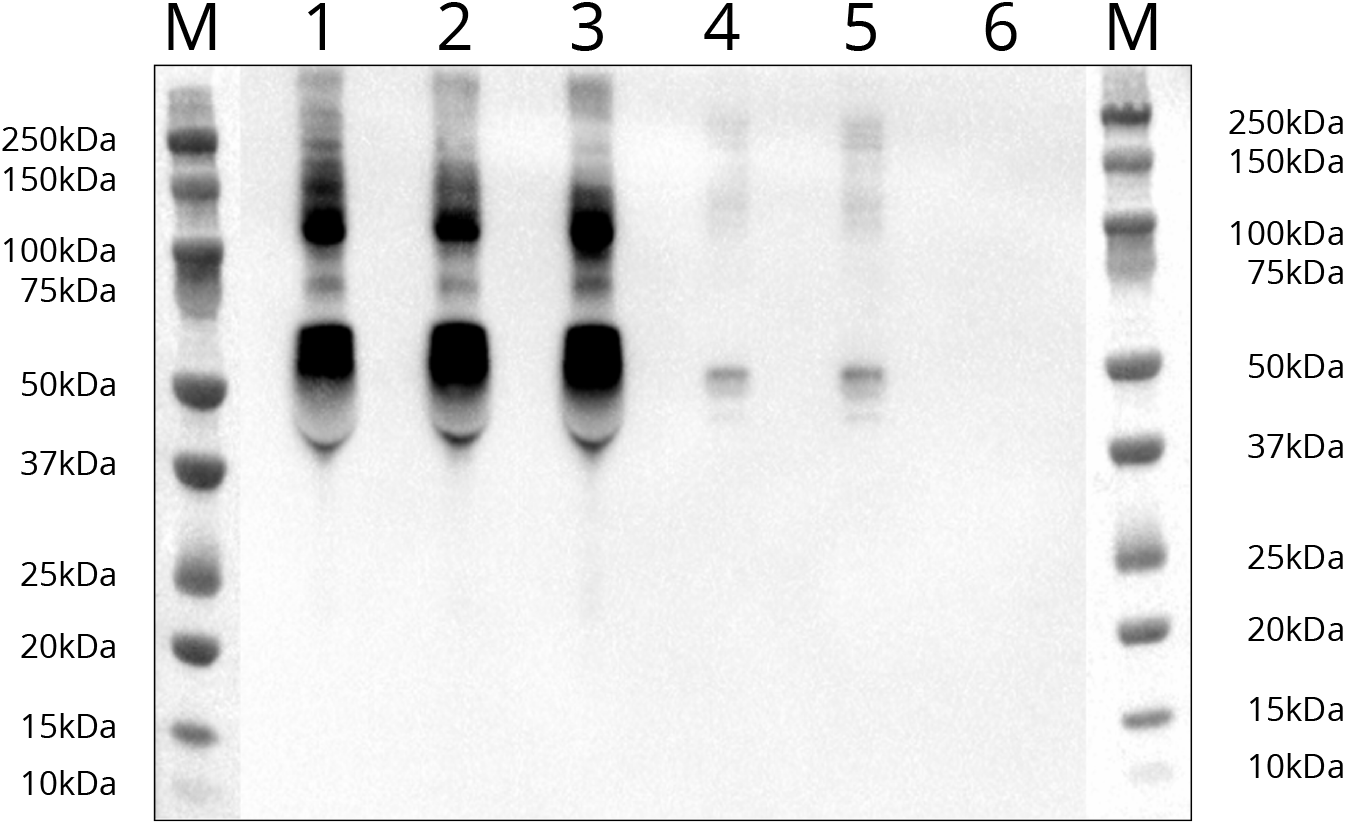
Western blot analysis of immunoreactive PMSG-associated signals in ELISA kit standards. Each sample was loaded in a 24 µL volume at a final concentration of 4 mIU/mL. Western blotting and detection were performed as described in Materials and Methods. M indicates marker (dual-colour BioRad); standards are as follows: 1) DRG, 2) Creative Diagnostics, 3) Demeditec, 4) Biomatik, 5) MyBiosource and 6) Krishgen.

**Figure 2:**
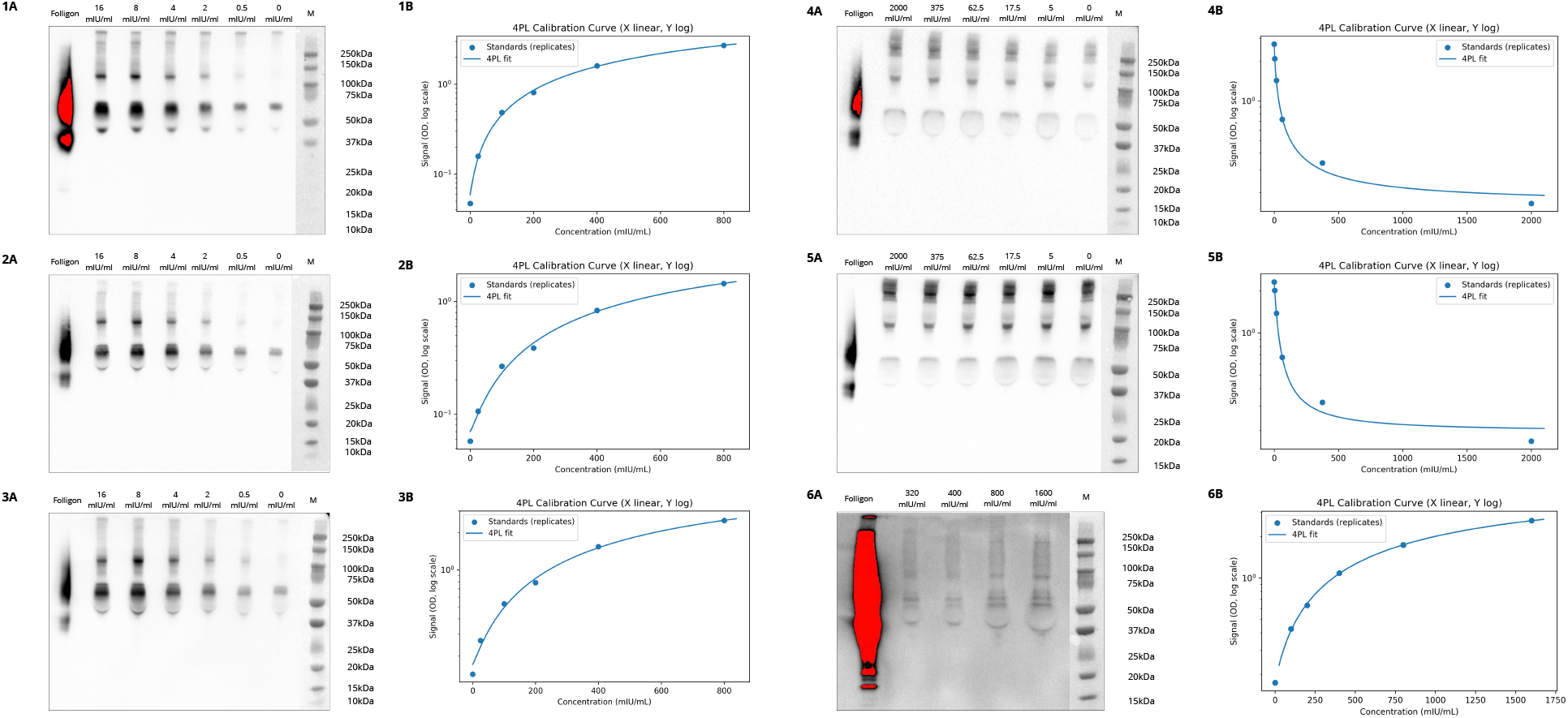
Comparison of kit-provided calibration series by Western blotting and ELISA standard-curve analysis. Kit-provided standards were analysed by Western blotting (A panels) and used to generate standard curves according to the respective manufacturer’s protocol using four-parameter logistic regression (B panels). For the DRG, Creative Diagnostics, and Demeditec Western blots, standards were diluted 1:50 before loading; consequently, the concentrations indicated above the lanes (16, 8, 4, 2, 0.5, and 0 mIU/mL) represent the final post-dilution concentrations loaded onto the gel, corresponding to original kit-standard concentrations of 800, 400, 200, 100, 25, and 0 mIU/mL. Biomatik, MyBiosource, and Krishgen standards were not subjected to this additional 1:50 dilution. Numbers indicate commercially marketed kits as follows: (1) DRG, (2) Creative Diagnostics, (3) Demeditec, (4) Biomatik, (5) MyBiosource, and (6) Krishgen.

No conversion from activity units to protein mass was performed. Where indicated, standards were adjusted to comparable nominal activity concentrations and loaded in equal volumes after dilution. Samples were resolved on a 4–20% gradient SurePAGE™ Bis-Tris gel (GenScript, M42015) under non-reducing conditions and transferred to a nitrocellulose membrane at 25 V and 125 mA for 90 min. Membrane was blocked in 5% milk in PBS-T for 1 h at room temperature and incubated with rabbit anti-PMSG antibody (1:1000) followed by HRP-conjugated anti-rabbit IgG antibody (1:2000, Cell Signaling Technology, cat. No. 7074S). Detection was carried out using chemiluminescence (SuperSignal™ West Pico Plus) and visualized with BioRad ChemiDoc.

### Coomassie staining

For Coomassie staining, the same sample preparation and electrophoresis conditions were used as described for Western blot analysis. Briefly, kit-provided standards were diluted in 0.9% NaCl to the indicated nominal concentrations and prepared under non-reducing conditions. Samples were resolved on a 4–20% gradient SurePAGE™ Bis-Tris gel (GenScript, M42015) under non-reducing conditions. After electrophoresis, the gel was stained with Coomassie Brilliant Blue staining solution and subsequently washed/destained overnight to reduce background staining and visualize the total protein pattern. Visualization was performed with BioRad ChemiDoc.

### Mass spectrometry

Protein standards provided with the respective ELISA kits were subjected to mass spectrometric analysis. Samples were analysed at the Functional Genomics Center Zurich (FGCZ, Zurich, Switzerland).

#### Sample preparation

##### In-Gel digestion

The gel pieces were washed three times with deionized water. They were destained three times with 50% acetonitrile in 25 mM ammonium bicarbonate (pH 8.5), followed by 100% acetonitrile. After evaporation of the organic solvent, proteins were digested with 0.5 µg of RapiZyme (Waters) at 60 °C for 30 min. The digested peptides were extracted by adding 50% acetonitrile with 0.1% TFA and sonicating for 10 min in a water bath. The supernatant was collected and dried by SpeedVac.

##### In-Solution digestion

For each product, 10 µg of protein (total protein concentration measured with UV/Vis polychromatic spectrophotometer Lunatic) was used for analysis. Proteins were reduced and alkylated in a solution containing 5 mM tris(2-carboxyethyl)phosphine and 15 mM chloroacetamide at 30 °C for 30 min. Proteins were then digested with 0.5 µg of RapiZyme (Waters) at 60 °C for 30 min. The samples were dried by SpeedVac.

##### LC-MS/MS analysis

Samples were analysed on a calibrated Q Exactive mass spectrometer (Thermo Fischer Scientific) coupled to a nano-Acquity UPLC system (Waters). Peptides were resuspended in 3% acetonitrile with 0.1% formic acid and loaded onto a nanoEase M/Z Symmetry (C18, 100A, 5 um, 1/PK, 180 um x 20 mm) and separated on a nanoEase M/Z HSS C18 Column (T3, 100A, 1.8 um, 1 P/K, 75 um x 250 mm), at a constant flow rate of 300 nL min-1, with a column temperature of 50°C and a linear gradient of 5−35% acetonitrile/0.1% formic acid in 45 min, and then 35-60% acetonitrile/0.1% formic acid in 5 min, followed by an increase to 95% acetonitrile in 10 min and then held isocratically for another 10 min. The mass spectrometer was operated in data-dependent acquisition (DDA) mode. One scan cycle comprised a full-scan MS survey spectrum followed by up to 12 sequential HCD MS/MS scans of the most intense precursors. Full-scan MS spectra (350–1500 m/z) were acquired in the FT-Orbitrap at a resolution of 70,000 at 400 m/z, while HCD MS/MS spectra were recorded in the FT-Orbitrap at a resolution of 30,000 at 400 m/z. HCD was performed with a target value of 1e5 and a normalised collision energy of 25 was applied. AGC target values were 1e5 for full FTMS. For all experiments, dynamic exclusion was used with a 10 s exclusion duration.

##### Data analysis

All LC-MS/MS raw data were searched against the UniProt horse database using the PEAKS Studio 10.5 (Bioinformatics Solutions Inc.) search engine, considering methionine oxidation and cysteine carbamidomethylation (for solution samples). The mzIdentML files were converted to Scaffold files (Proteome Software Inc.). Data are available via ProteomeXchange with identifier PXD075575.

## Results

To characterize the immunoreactive profile and apparent molecular-weight distribution of the kit-provided PMSG standards, we performed Western blot analysis using a polyclonal anti-PMSG (OriGene, AP02036SU-N) antibody (Figure 1). The highest-concentration standard supplied with each kit was diluted with 0.9% NaCl to a common nominal concentration of 4 mIU/mL before loading, allowing comparison of immunoreactive profiles at the same nominal activity concentration. Standards from three kits (DRG, CD, and Demeditec) obtained from different suppliers produced highly similar banding patterns. Two additional kits (Biomatik and MyBiosource) also showed highly similar patterns to each other, but with weaker signals than the first three kits. Under the non-reducing conditions used in this study, PMSG/eCG was expected to appear predominantly within a broad apparent molecular-weight range of approximately 45–70 kDa, reflecting its highly glycosylated heterodimeric structure. Consistent with this expectation, five of the six kit standards showed a dominant immunoreactive signal between 45 and 55 kDa, with a broader signal extending within the expected molecular-weight range. In addition, higher-molecular-weight immunoreactive bands were observed, most prominently around 100 kDa and around 75, 150, and 160 kDa (Figure 1). No bands corresponding to single-chain forms were detected. The Krishgen standard did not produce a detectable immunoreactive signal under these conditions. To assess total protein loading and overall protein composition independently of antibody-based detection, the same kit standards loaded at 4 mIU/mL were analysed on a parallel Coomassie-stained SDS–PAGE gel (Supplementary Figure 1). Coomassie staining showed clear differences in total protein abundance and banding patterns among standards loaded at the same nominal PMSG concentration. DRG, Creative Diagnostics, and Demeditec produced similarly intense, complex profiles, whereas Biomatik and MyBiosource showed weaker and less complex patterns. The Krishgen standard produced only a faint band at approximately 50 kDa.

To evaluate the internal consistency and quantitative validity of the ELISA kit standards, serial dilutions of each manufacturer-provided standard were analysed independently by Western blot (Figure 2 a-panel) and in parallel used to generate the corresponding standard curves according to the manufacturers’ protocols (Figure 2 b-panel). Samples for Western blotting were loaded based on the declared concentrations and processed under identical experimental conditions. Due to the high protein concentrations of the standards provided by DRG, Creative Diagnostics, and Demeditec, samples were diluted 1:50 for the final Western blot analysis to improve visualization. In the first tested kits (Figure 2, 1-3) the highest standard concentration reflects the strongest band, and a progressive dilution shows a stepwise decrease in band intensity.

In a competitive ELISA, signal intensity in the assay decreases with increasing analyte concentration, which we can see in Figure 2 (4b and 5b). However, this competitive behaviour occurs within the ELISA plate format, not in a Western blot. Western blot detects absolute protein amounts and does not reflect competitive binding dynamics. Therefore, band intensity should increase with increasing protein concentration, regardless of the direction of the ELISA signal. However, standards from Biomatik, MyBiosource, and Krishgen lacked a dose-response effect and showed consistent band intensity regardless of concentration. The zero calibrators, which had a nominal PMSG concentration of 0 mIU/mL, produced a detectable immunoreactive signal in the same apparent molecular-weight range as the non-zero standart (Figure 2). The signal intensity was comparable to that observed for several higher nominal standards. To determine whether the Western blot signal in the zero standards was associated with detectable protein material in the calibrator matrix, the corresponding zero-standard preparations were analysed on a parallel Coomassie-stained SDS–PAGE gel (Supplementary Figure 2). Coomassie staining revealed visible protein bands in several nominal zero-standard lanes, indicating that these calibrators were not protein-free matrices despite being supplied as zero or target-free standards.

Following the non-uniform Western blot signals observed among the different ELISA kit standards when probed with a polyclonal anti-PMSG antibody, we sought to determine whether these differences reflected variations in protein composition rather than antibody-binding heterogeneity. Given that polyclonal antibodies recognize multiple epitopes and may detect isoforms or related proteins, we performed an independent analytical approach to directly assess the molecular composition of the standards. To this end, all kit-provided standards were normalised to identical total protein concentrations and subjected to mass spectrometry analysis. Figure 3 shows a heatmap that summarizes the exclusive unique peptide counts (log scale) detected for each protein in standards. Protein profiling revealed that standards from DRG, CD and Demeditec exhibited highly similar protein compositions, as did standards from MyBiosource and Biomatik. In contrast, the Krishgen standard showed the lowest overall protein content among all analysed samples, which corresponded well with the Western blot results. Notably, none of the tested preparations contained the protein of interest or closely related members of the same protein family. Serum albumin was detected in all analysed standards and showed the highest peptide counts across suppliers. Several additional plasma-associated proteins were consistently identified in multiple kits (DRG, Creative Diagnostics, Demeditec, MyBiosource, and Biomatik), including serotransferrin, haptoglobin, hemopexin, ceruloplasmin, and fibronectin. These proteins have distinct physiological functions: serotransferrin and ceruloplasmin are involved in metal-ion transport and homeostasis; haptoglobin and hemopexin bind haemoglobin and free haem, respectively; and fibronectin is an extracellular-matrix and plasma glycoprotein involved in cell adhesion, tissue repair, and coagulation-related processes. Apolipoproteins (ApoA1, ApoA2, ApoA4, ApoB, ApoC2, and ApoH) were identified mainly in DRG, Creative Diagnostics, and Demeditec, indicating the presence of lipoprotein-associated plasma fractions in these preparations. Complement C5 and Complement C9, components of the complement cascade, were detected in MyBiosource and Biomatik preparations. Krishgen standards were characterised by a markedly reduced protein diversity compared to other suppliers, with serum albumin representing the dominant identified protein. Overall, most proteins identified in the kit standards were plasma-associated rather than specific to PMSG.

**Figure 3:**
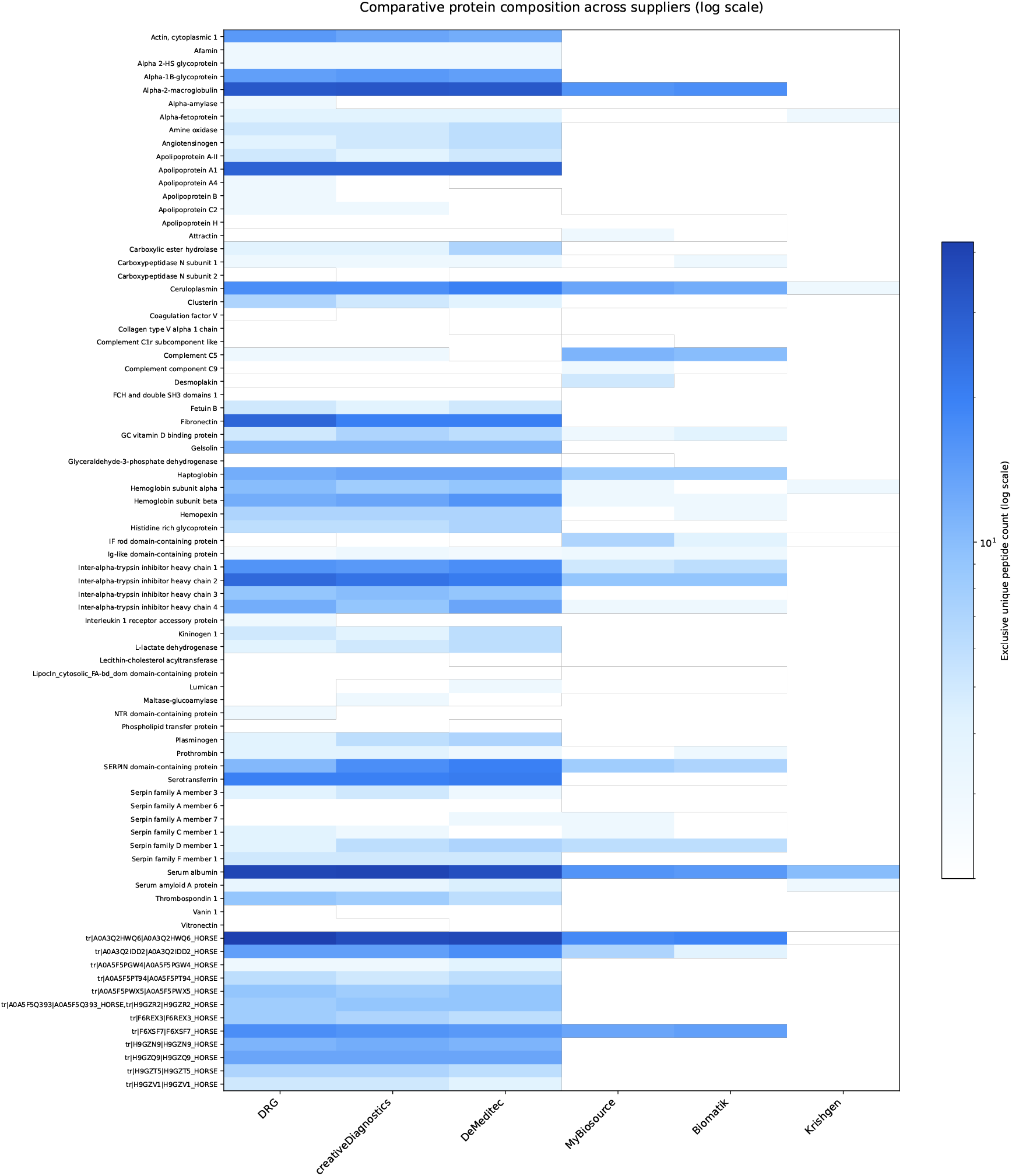
Comparative protein composition of ELISA kit standards across suppliers. Heatmap showing exclusive unique peptide counts (log scale) obtained by mass spectrometry for proteins identified in kit-provided standard solutions at the maximum concentration of each kit from DRG (800 mIU/mL), Creative Diagnostics (800 mIU/mL), Demeditec (800 mIU/mL), MyBiosource (2000 mIU/mL), Biomatik (2000 mIU/mL), and Krishgen (1600 mIU/mL). Rows represent identified proteins; columns represent suppliers. Colour intensity corresponds to the log-transformed number of exclusive unique peptides detected per protein in each solution.

To compare quantitative performance across suppliers, commercial PMSG preparations with known nominal concentrations were serially diluted and measured using ELISA kits from six manufacturers. Measured concentrations were plotted against nominal concentrations for each kit (Figure 4). Quantitative regression analysis confirmed distinct differences in concentration-dependent assay behaviour. DRG, Creative Diagnostics, and Demeditec showed clear positive relationships between nominal and measured concentrations for both Folligon and OPPA, with regression slopes ranging from 0.519 to 0.778 and *R*^2^ values from 0.897 to 1.000. However, all slopes remained below the theoretical value of 1, and the mean absolute deviation from nominal concentrations ranged from 34.9% to 46.5%, indicating systematic quantitative disagreement despite the concentration-dependent response. In contrast, Biomatik and MyBiosource showed regression slopes close to zero and mean absolute deviations of approximately 99%, consistent with the absence of a meaningful concentration-dependent quantitative response. The Krishgen assay similarly showed low slopes (0.053–0.059) and *R*^2^ values of 0.313–0.450, reflecting the relatively narrow range of measured concentrations despite substantial changes in nominal input. Detailed regression parameters are provided in Supplementary Table 2.

**Figure 4:**
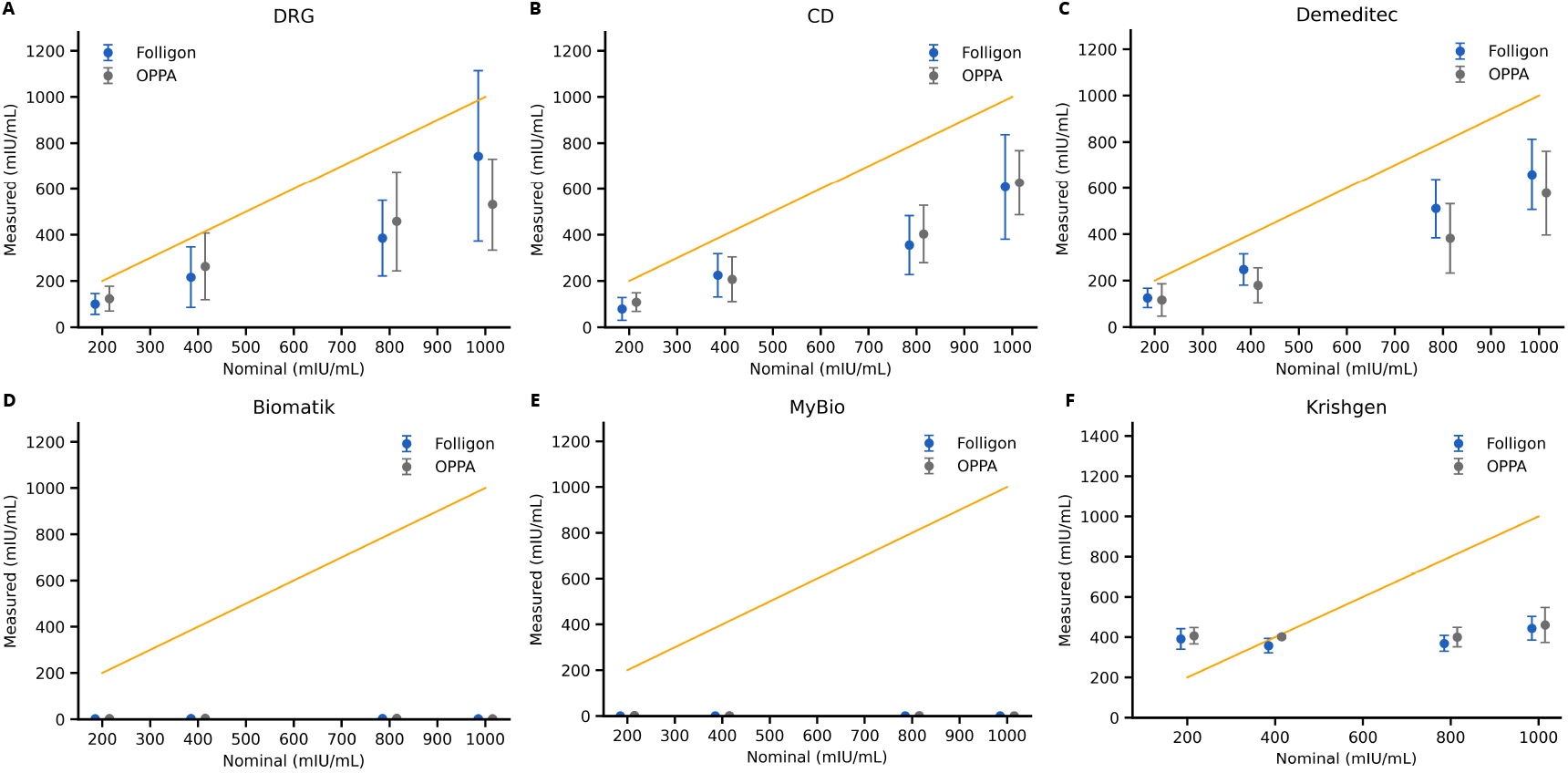
Comparison of measured versus nominal PMSG concentrations across ELISA kits from different suppliers. Commercial PMSG preparations (Folligon and OPPA) were serially diluted to defined nominal concentrations and measured using ELISA kits from six suppliers: (A) DRG, (B) Creative Diagnostics (CD), (C) Demeditec, (D) Biomatik, (E) MyBiosource (MyBio), and (F) Krishgen. Measured concentrations (mIU/mL) are plotted against nominal concentrations (mIU/mL). The orange line indicates the theoretical 1:1 relationship between nominal and measured values. Data points represent mean ± SD. n = 3 (except Krishgen)

## Discussion

Several commercially available ELISA kits marketed as quantitative were evaluated to determine their suitability for PMSG measurement (Table 1). Across assays, we observed limited ability to reliably distinguish between different nominal concentrations. In particular, standard curve behaviour and sample dilution responses did not consistently demonstrate proportional signal changes across the expected working range.

ELISA detects immunoreactivity through antibody recognition of accessible epitopes rather than measuring protein mass or biological activity. For PMSG, this distinction is relevant because glycosylation and molecular heterogeneity may affect antibody binding. Native PMSG isolated from equine serum exhibits extensive and variable glycosylation [13], including differences in branching and sialylation [4]. These structural features can influence epitope accessibility [11] and antibody binding, potentially leading to preferential detection of specific molecular forms [14]. Consequently, quantitative consistency depends critically on the nature and composition of the reference standard used for calibration. This raises the possibility that inconsistencies in ELISA measurements may originate not only from assay design but also from the molecular composition of the calibration standards.

To further investigate these inconsistencies, we analysed the composition of kit-provided standards using Western blot and mass spectrometry. To assess the authenticity and purity of ELISA kit standards purported to contain PMSG, Western blot analysis was performed using a polyclonal anti-PMSG antibody (Figure 1). Western blot analysis revealed marked differences in banding patterns between the evaluated ELISA kit standards. Several kits produced a dominant signal within the expected apparent molecular-weight range for highly glycosylated PMSG/eCG, whereas additional higher-molecular-weight immunoreactive bands were also observed. The identity of the additional immunoreactive bands could not be determined by Western blotting alone. Their presence shows that the antibody detected several molecular species rather than a single band within the expected PMSG range.

Notably, the “zero standard” supplied by five out of six tested manufacturers - designated as a negative control and reported to be devoid of target protein - produced a clear detectable immunoreactive band at the expected molecular weight (Figure 2). The signal intensity was comparable to several higher nominal standards. Although a polyclonal antibody was used for detection, which may detect related proteins or fragments, the consistent detection of a band of appropriate size in the zero standard cannot readily be explained by non-specific cross-reactivity alone. Parallel Coomassie staining of the same zero-standard preparations further showed that these calibrators contained detectable total protein (Supplementary Figure 2). This finding supports the interpretation that the zero standards represent protein-containing matrices rather than truly protein-free controls. The presence of immunoreactive signal in nominally protein-free standards raises additional concerns regarding the definition and quality control of calibration materials.

Mass spectrometric profiling further indicated discrepancies between the declared and actual protein composition of the kit standards. In addition to the analysis of total standard solutions, we performed an orthogonal verification experiment to address the possibility that low-abundance PMSG might have escaped detection in bulk protein mixtures. Standards were separated by SDS– PAGE in parallel to Western blotting and stained with Coomassie (Supplementary Figure 3).

Bands corresponding to the expected molecular weight range were excised and subjected to in-gel digestion followed by LC–MS/MS analysis (Data are available via ProteomeXchange with identifier PXD075575). Despite targeted analysis of gel regions matching the immunoreactive signal, PMSG-derived peptides were not identified. This observation further supports the conclusion that the immunoreactive bands detected by Western blot do not correspond to a purified or dominant PMSG species within the analysed standards. Taken together, the discordance between Western blot detection, ELISA calibration curve behaviour, and orthogonal protein identification suggests inconsistencies in standard composition that may affect quantitative reliability. Importantly, these findings raise concerns regarding the commutability of kit-provided standards. In quantitative immunoassays, calibration material must behave in a manner comparable to the analyte present in real samples across dilution ranges and assay conditions. If the molecular composition of the standard differs substantially from that of the target preparation, interpolation against the calibration curve may not reflect the true immunoreactive content of the sample. For structurally heterogeneous glycoproteins such as PMSG, even subtle compositional differences between standards may translate into disproportionate effects on signal proportionality. The observed lack of parallel dilution behaviour in several kits suggests that this requirement may not be consistently fulfilled.

Notably, the kits that demonstrated proportional signal behaviour across nominal dilution ranges were also those exhibiting comparable protein profiles and consistent dilution-dependent band intensities in Western blot analysis. In contrast, kits lacking dose-dependent ELISA responses showed either uniform band intensities across concentrations or markedly distinct protein compositions by mass spectrometry. This internal concordance across orthogonal analytical approaches suggests that quantitative performance is closely linked to the molecular characteristics of the respective calibration standards.

Western blotting provides information on immunoreactive bands and their apparent molecular weight, but it does not directly identify the proteins present in each band or provide a direct measure of sample purity [15]. LC–MS/MS provides protein-level identification, although highly glycosylated or low-abundance eCG may remain undetected in complex mixtures. In our previous study, both α- and β-subunits of eCG/eLH were identified by LC–MS/MS in the same commercial PMSG preparations and lots used here as reference preparations, but at a higher total protein input [4]. The absence of eCG-derived peptides from the kit standards should therefore be interpreted as non-detection under the conditions used [16]. ELISA results are also dependent on antibody recognition, calibration material, and assay format. For this reason, agreement between Western blot, mass spectrometry, and ELISA should be interpreted cautiously and should not be considered definitive molecular identification of each immunoreactive band. Nevertheless, the same general pattern was observed across the different analytical approaches. Standards with dilution-dependent Western blot signals also showed related protein profiles by mass spectrometry and more proportional ELISA responses. In contrast, standards with weak or non-proportional Western blot signals showed distinct protein compositions and limited proportionality in ELISA measurements. The related patterns observed by Western blotting, mass spectrometry, and ELISA show that the differences between kits were not confined to a single analytical method. However, the present experiments do not establish which properties of the calibration standards caused the differences in quantitative response.

Quantitative comparison of measured and nominal concentrations across ELISA kits further illustrated these inconsistencies (Figure 4). In principle, accurate quantitative assays should produce measurements that scale proportionally with the nominal concentration of the analyte, approximating the theoretical 1:1 relationship. While three assays (DRG, Creative Diagnostics, and Demeditec) showed a general increase in measured concentration with increasing nominal input, deviations from the theoretical relationship were still observed. In contrast, other kits demonstrated little or no proportional response across the tested dilution range. In these cases, measured concentrations remained close to baseline or changed only minimally despite substantial differences in nominal input. These observations indicate that the relationship between nominal and measured PMSG concentration is not consistently preserved across assays. In immunoassays, the measured signal may be influenced not only by the amount of target analyte, but also by the composition of the surrounding sample matrix [17, 18, 19]. Components such as carrier proteins, serum-derived proteins, buffer constituents, salts, preservatives, or other co-purified material may affect background signal, nonspecific binding, antibody-antigen interaction, and epitope accessibility. Therefore, samples containing comparable amounts of immunoreactive PMSG may generate different ELISA responses if their matrix differs from that of the kit calibrator. This is also relevant for PMSG preparations and kit-provided standards, which consist of complex serum-derived protein mixtures rather than purified, structurally uniform analytes [4]. In the present study, mass spectrometry revealed marked differences in the protein composition of kit standards, including serum albumin and other plasma-associated proteins. Thus, differences between nominal and measured concentrations may reflect not only differences in PMSG content or antibody recognition, but also matrix-dependent effects. The atypical behavior observed for the Krishgen assay, where measured concentrations remained within a relatively narrow range despite changes in nominal input, may be explained by matrix-dependent background signal, non-parallel behavior between the tested preparation and the kit calibrator, or differences in the composition of the calibration material. Dilution-parallelism and spike-and-recovery experiments would be needed to determine whether the restricted response of the Krishgen assay arose from matrix effects or from differences between the kit calibrator and the tested preparations [20, 21].

These findings are particularly relevant given the absence of a universally accepted, commutable reference preparation [22]. Following depletion of the WHO/Ph. Eur. standard and suspension of the eCG monograph, no broadly adopted calibration material is currently available. In practice, many PMSG preparations consist of complex equine serum-derived protein mixtures, in which PMSG co-exists with other serum proteins. Such matrices may introduce cross-reactivity, background signal, or altered epitope presentation, thereby complicating immunoassay-based quantification.

For an ELISA to be considered quantitatively valid, several criteria should be met within the relevant biological matrix, including proportional signal response, parallelism between diluted samples and standard curves, accurate recovery of spiked analyte, and acceptable intra- and inter-assay variability. For PMSG, fulfilment of these criteria is inherently challenging due to molecular heterogeneity, matrix effects, and the lack of a standardized reference material. An additional conceptual limitation arises from the use of International Units (IU) as the primary measure of PMSG potency. IU reflect biological activity, whereas ELISA measures epitope presence. Because the relationship between molecular mass and biological potency varies depending on glycosylation and structural form, immunoreactivity cannot be assumed to correlate directly with bioactivity without complementary functional validation. This conceptual discrepancy becomes particularly relevant for PMSG, as International Units are historically defined through *in vivo* bioassays that integrate receptor binding [12], signal transduction, and downstream physiological effects. In contrast, ELISA quantifies antibody-accessible epitopes without regard to receptor interaction or functional potency. Given that different glycoforms of PMSG may exhibit altered receptor affinity while retaining immunoreactive epitopes, the relationship between IU-based potency and ELISA-derived concentration cannot be presumed to be linear or constant across preparations. Consequently, calibration of immunoassays against IU-defined standards may introduce systematic deviations when structural heterogeneity differs between reference material and tested samples.

Based on the analytical limitations identified here, a future PMSG/eCG reference material should be comprehensively characterised using various analytical approaches. Protein identity and purity could be assessed by mass spectrometry-based peptide mapping in combination with chromatographic methods such as RP-HPLC or SEC, while molecular heterogeneity could be evaluated by intact-mass analysis, size-exclusion and/or charge-based separation techniques. Glycosylation should be characterised using glycan profiling and, where appropriate, site-specific glycopeptide analysis by LC-MS/MS. In addition to this physicochemical characterisation, biological potency should be independently assigned using a validated functional bioassay, preferably one capable of assessing the relevant FSH- and/or LH-receptor activity of eCG. Such an integrated approach would provide both structural and functional characterisation of the reference material and would reduce the uncertainty associated with assigning its biological activity based solely on physicochemical measurements. Development and formal validation of such a standard, however, are beyond the scope of the present Technical Note.

Collectively, our findings indicate that currently available commercial ELISA kits may exhibit limitations in quantitative accuracy when applied to heterogeneous PMSG preparations. Systematic validation against well-characterised reference material, or the development of standardized calibration preparations, will be essential to improve repeatability, inter-laboratory comparability, and interpretability of quantitative PMSG measurements in both research and veterinary contexts.

## Conclusion

Reliable quantitative determination of equine chorionic gonadotropin (eCG/PMSG) by ELISA remains challenging because current commercial assays do not consistently measure PMSG preparations in a proportional and comparable manner. In this study, different kits produced substantially different quantitative outputs for the same nominal PMSG concentrations, suggesting that assay format, antibody specificity, calibration material, matrix composition, and the fact that assay calibration is commonly expressed in International Units (IU) all contribute to inter-assay variability. These limitations are further amplified by PMSG structural heterogeneity and variable glycosylation, which may alter epitope accessibility and reduce immunological comparability between calibrators and test samples. Thus, the absence of a validated, commutable international reference material remains a major barrier to standardisation alongside uncertain immunological comparability between kit calibrators and tested PMSG preparations. Therefore, this would be an important step toward standardisation and inter-laboratory comparability, but reliable quantification will also require assay-specific validation, including dilution parallelism and recovery testing in relevant sample matrices.

## Acknowledgements

We would like to thank Zdenka Herceg for her attention to detail and for pointing out these difficulties during her internship. We would also like to thank Hafida Atiqui, who supported us throughout with organizational work related to ordering and shipment handling, making this comparison possible. We also thank Dr. Mai Azuma, who helped shape this paper with insightful questions. We also thank Dr. Chia-Wei Lin for support with Mass spectrometry experiments.

