## Supplementary Data for "Standardisation Challenges in ELISA-Based Quantification of Equine Chorionic Gonadotropin (eCG/PMSG)"

**Supplementary Table 1.** Overview of used ELISA kits and protocol comparison.
**Supplementary Table 2.** Quantitative comparison of nominal and ELISA-measured PMSG concentrations.
**Supplementary Figure 1.** Coomassie gel of the 4 mIU/mL sample corresponding to main Figure 1.
**Supplementary Figure 2.** Coomassie gel of undiluted zero-standard samples.
**Supplementary Figure 3.** Comparison of Western blot and Coomassie gels analysed by mass spectrometry.

Supplementary Table 1: Overview of used ELISA Kits and Protocol comparison

| **ELISA kit** | **DRG** | **Creative Diagnostics** | **Demeditec** | **MyBiosource** | **Biomatik** | **Krishgen** |
|---|---|---|---|---|---|---|
| Lot # | 17K095-2 | 17K104 | 17K055 | C1465081439 | C1478041497 | EPMSG0425 |
| Ref./Cat.# | EIA-1298 | DEIA2235 | DE1298 | MBS701410 | EKC38379 | KLV0061 |
| Exp. Date (DD.MM.YYYY) | 30.09.2026 | 31.10.2025 | 31.05.2026 | 19.09.2025 | 27.02.2026 | 31.03.2026 |
| Ordered / Arrived | Mai 2025 / October 2025 | March 2025 / | March 2025 / | March 2025 / | August 2025 / | March 2025 / |
| Components | Microtiter 96 Wells (12x8) : Wells coated with Anti PMSG (monoclonal) Lot: 17W095 Enzyme conjugate (11ml) : Anti-PMSG antibody conjugated with horseradish peroxidase Lot: 17D095 Substrate Solution (14ml) : Tetramethylbenzidine (TMB) Lot: TMBS250007 Stop solution (14ml): 0.5M H2SO4 Lot: SD085 Standards (6x0.5ml) : Standards are calibrated against WHO international Standard Serum Gonadotropin, Equine (NIBSC 62/001) Lot: 17S095 Wash solution 40x (30ml) Lot: TWP065 Assay buffer (25ml) Lot: 17PB095 | Microtiter 96 Wells (12x8): Wells coated with Anti PMSG (monoclonal) Enzyme conjugate (11ml): Anti-PMSG antibody conjugated with horseradish peroxidase Substrate Solution (14ml): Tetramethylbenzidine (TMB) Stop solution (14ml): 0.5M H2SO4 Standards (6x0.5ml): Standards are calibrated against WHO international Standard Serum Gonadotropin, Equine (NIBSC 62/001) Wash solution 40x (30ml) Assay buffer (25ml) | Microtiter 96 Wells (12x8) : Wells coated with Anti PMSG (monoclonal) Lot: 17W035 Enzyme conjugate (11ml) : Anti-PMSG antibody conjugated with horseradish peroxidase Lot: 17D055 Substrate Solution (14ml) : Tetramethylbenzidine (TMB) Lot: TMBS240013 Stop solution (14ml): 0.5M H2SO4 Lot: SD025 Standards (6x0.5ml) : Standards are calibrated against WHO international Standard Serum Gonadotroping, Equine (NIBSC 62/001) Lot: 17S054 Wash solution 40x (30ml) Lot: TWP045 Assay buffer (25ml) Lot: 17PB045 | Assay plate 96 Wells: Precoated with goat-anti-rabbit antibody Lot: CAP25146508 Standards (6x0.5ml): Lot: CS25146508 Antibody (6ml): Against PMSG (Horse (Equus caballus; Equine) and HRP-labelled PMSG Lot: CA25146508 horseradish peroxidase (HRP)-conjugated (6ml) Lot: CHC25146508 Wash buffer 20x (15ml) Lot: CWB25146508 Substrate A (7ml) Lot: CSA25146508 Substrate B (7ml) Lot: CSB25146508 Stop solution (7ml) Lot: CSS25146508 Adhesive strip for 96 Well plate x4 | Assay plate 96 Wells: Precoated with goat-anti-rabbit antibody Lot: CAP25147804 Standards (6x0.5ml): Lot: CS25147804 Antibody (6ml): Against PMSG (Horse (Equus caballus; Equine) and HRP-labelled PMSG Lot: CA25147804 horseradish peroxidase (HRP)-conjugated (6ml) Lot: CHC25147804 Wash buffer 20x (15ml) Lot: CWB25147804 Substrate A (7ml) Lot: CSA25147804 Substrate B (7ml) Lot: CSB25147804 Stop solution (7ml) Lot: CSS25147804 Adhesive strip for 96 Well plate x4 | Equine PMSG Antibody (monoclonal) coated Microtiter Plate (8x12 wells) Lot: EPMSG0425 Standard, Equine PMSG (0.5ml): concentrated Lot: EPMSG0425 Biotinylated PMSG Antibody (1ml) Lot: EPMSG0425 Streptavidin HRP-conjugate (6ml) Lot: EPMSG0425 Standard diluent (3ml) Lot: EPMSG0425 Sample diluent 1x (12ml) Lot: EPMSG0425 Wash buffer 20x (25ml) Lot: EPMSG0425 Tetramethylbenzidine (TMB) substrate (12ml) Lot: EPMSG0425 Stop solution (12ml) Lot: EPMSG0425 |

Continued on next page

Supplementary Table 1: Overview of used ELISA Kits and Protocol comparison (continued)

| **ELISA kit** | **DRG** | **Creative Diagnostics** | **Demeditec** | **MyBiosource** | **Biomatik** | **Krishgen** |
|---|---|---|---|---|---|---|
| Standard concentrations | S0: 0 mIU/ml; S1: 25 mIU/ml; S2: 100 mIU/ml; S3: 200 mIU/ml; S4: 400 mIU/ml; S5: 800 mIU/ml; | S0: 0 mIU/ml; S1: 25 mIU/ml; S2: 100 mIU/ml; S3: 200 mIU/ml; S4: 400 mIU/ml; S5: 800 mIU/ml; | S0: 0 mIU/ml; S1: 25 mIU/ml; S2: 100 mIU/ml; S3: 200 mIU/ml; S4: 400 mIU/ml; S5: 800 mIU/ml; | S0: 0 mIU/ml; S1: 5 mIU/ml; S2: 17.5 mIU/ml; S3: 62.5 mIU/ml; S4: 375 mIU/ml; S5: 2000 mIU/ml; | S0: 0 mIU/ml; S1: 5 mIU/ml; S2: 17.5 mIU/ml; S3: 62.5 mIU/ml; S4: 375 mIU/ml; S5: 2000 mIU/ml; | Concentrated standards: 3200 mIU/ml |
| Protocol version | Version 6.0 2021-08-20 | 21-04/25 | 06-08/21/DMC Updated 210823 | Protocol from Catalog number: MBS701410 Version not available | Protocol from Catalog number: EKC38379 Version not available | Protocol from Catalog number: KLV0061 Version 1.0 |

Supplementary Table 2: Quantitative comparison of nominal and ELISA-measured PMSG concentrations.

| **ELISA kit** | **PMSG preparation** | **Slope** | **Intercept (mIU/mL)** | $R^2$ | **Mean absolute relative deviation (%)** |
|---|---|---|---|---|---|
| DRG | Folligon | 0.778 | -76.7 | 0.897 | 38.6 |
| | OPPA | 0.519 | 55.3 | 0.976 | 36.1 |
| Creative Diagnostics | Folligon | 0.621 | -37.7 | 0.938 | 46.5 |
| | OPPA | 0.622 | -23.1 | 0.970 | 42.3 |
| Demeditec | Folligon | 0.659 | -4.0 | 1.000 | 34.9 |
| | OPPA | 0.585 | -18.6 | 0.967 | 44.3 |
| Biomatik | Folligon | 0.0001 | 2.59 | 0.060 | 99.4 |
| | OPPA | -0.0007 | 3.50 | 0.109 | 99.2 |
| MyBiosource | Folligon | 0.0002 | 0.70 | 0.400 | 99.8 |
| | OPPA | -0.0018 | 2.83 | 0.776 | 99.5 |
| Krishgen | Folligon | 0.059 | 355.4 | 0.313 | 53.9 |
| | OPPA | 0.053 | 385.7 | 0.450 | 52.0 |

$T$echnical replicates were averaged within each independent ELISA experiment before analysis. Regression parameters were obtained by ordinary least-squares regression of the mean measured concentration against the corresponding nominal concentration. A slope of 1 and an intercept of 0 represent theoretical perfect agreement.

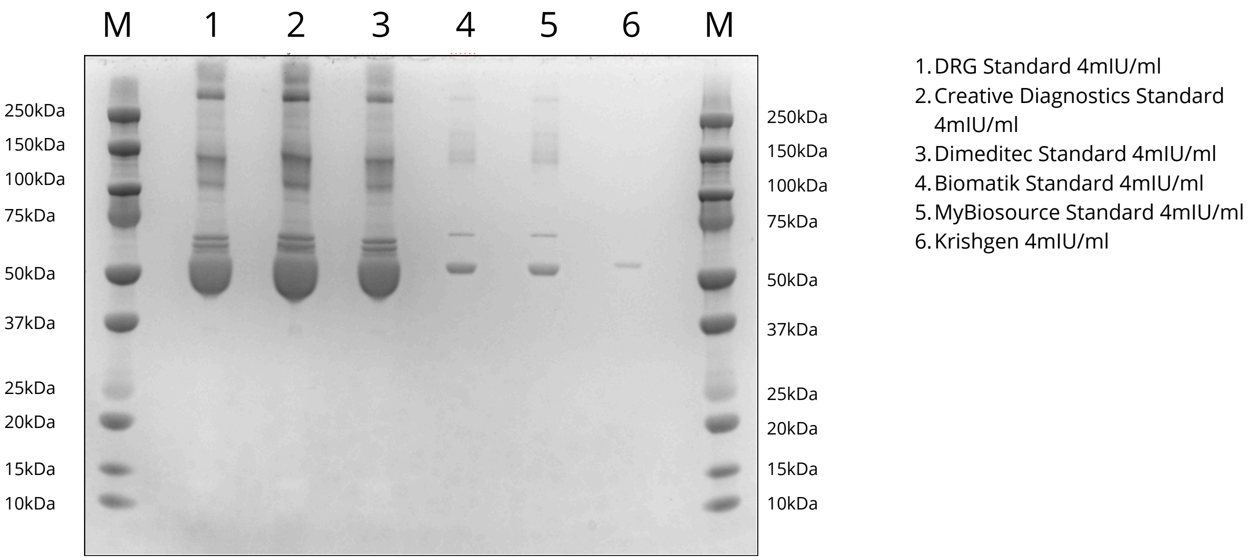


Supplementary Figure 1: Coomassie Gel of standards provided by each Kit diluted up to 4mIU/ml, loaded in non-reduced state. M indicates Marker (dual-colour BioRad); standards are as follows: 1) DRG, 2) Creative Diagnostics, 3) Dimeditec, 4) Biomatik, 5) MyBiosource and 6) Krishgen.

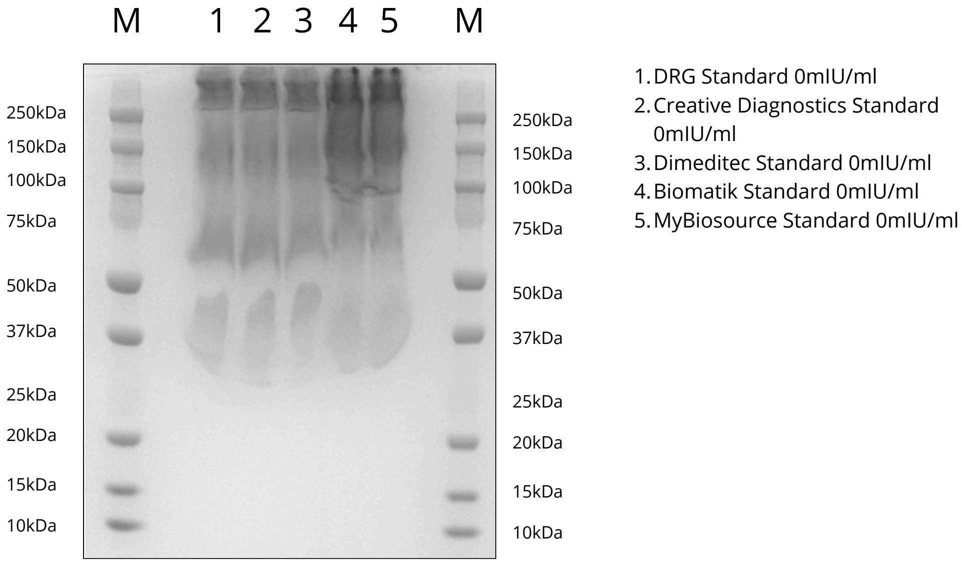


Supplementary Figure 2: Coomassie Gel of zero standards provided by each Kit, loaded in non-reduced state. M indicates Marker (dual-colour BioRad); standards are as follows: 1) DRG, 2) Creative Diagnostics, 3) Dimeditec, 4) Biomatik, 5) MyBiosourc.

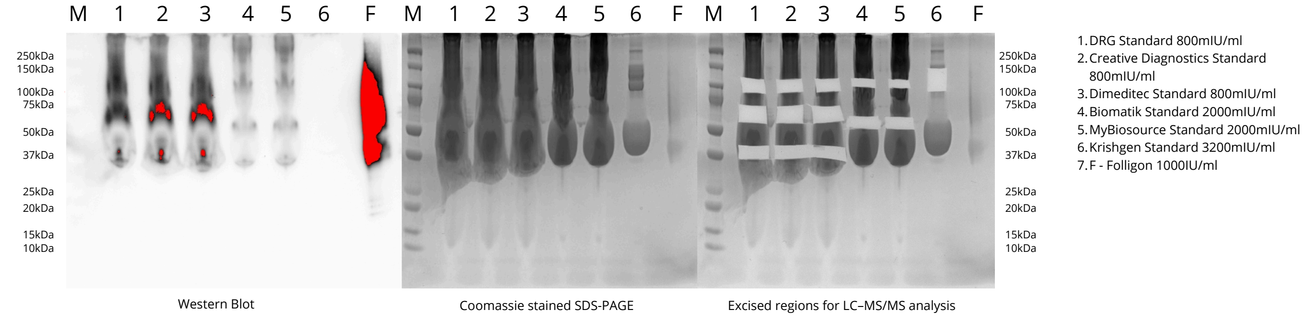


Supplementary Figure 3: Western blot analysis, parallel Coomassie-stained SDS-PAGE gel, and gel regions excised for LC–MS/MS analysis based on comparison with the Western blot signal. For each sample, multiple excised gel bands were pooled and analyzed together as a single LC–MS/MS sample. Samples loaded in non-reduced state in maximum concentration of each kit. M indicates Marker (dual-colour BioRad), F indicates Folligon; standards are as follows: 1) DRG, 2) Creative Diagnostics, 3) Dimeditec, 4) Biomatik, 5) MyBiosource and 6) Krishgen.
